# Single-Nucleus RNA Sequencing of Developing and Mature Superior Colliculus Identifies Neuronal Diversity and Candidate Mediators of Circuit Assembly

**DOI:** 10.1101/2023.02.01.526254

**Authors:** Ana C. Ayupe, James S. Choi, Felipe Beckedorff, Paola Catanuto, Robyn Mccartan, Konstantin Levay, Kevin K. Park

## Abstract

The superior colliculus (SC) is a sensorimotor structure in the midbrain that integrates input from multiple sensory modalities to initiate motor commands. It undergoes well-characterized steps of circuit assembly during development, rendering the mouse SC a popular model to study establishment and refinement of neural connectivity. Here we performed single nucleus RNA-sequencing analysis of the mouse SC isolated at various developmental time points. Our study provides a transcriptomic landscape of the cell types that comprise the SC across murine development with particular emphasis on neuronal heterogeneity. We used these data to identify Pax7 as a marker for an anatomically homogeneous population of GABAergic neurons. Lastly, we report a repertoire of genes differentially expressed across the different postnatal ages, many of which are known to regulate axon guidance and synapse formation. Our data provide a valuable resource for interrogating the mechanisms of circuit development, and identifying markers for manipulating specific SC neuronal populations and circuits.

## INTRODUCTION

The superior colliculus (SC) is a paired sensorimotor structure in the midbrain that receives input from multiple sensory modalities and incorporates environmental stimuli to control innate behaviors. In the mouse, these behaviors include coordinating gaze shifts involving both eye and head movements (Sparks and Mays, 1990), escaping or freezing in response to a looming object (Shang et al., 2018; Wei et al., 2015), hunting and approaching responses (Shang et al., 2019), and light-/dark-induced sleep and wakefulness (Zhang et al., 2019).

The SC can be divided into the superficial visuosensory and the deeper motor layers with each subdivision organized into several fibrous and cellular laminae (Basso and May, 2017; Benavidez et al., 2021; Cang et al., 2018; Ito and Feldheim, 2018; May, 2006). The superficial region consists of the stratum griseum superficiale (SGS) and the stratum opticum (SO) and receives innervation from retinal ganglion cells (RGCs) and the primary visual cortex. Cells in the deeper layer express sensitivity to sensory stimuli from multiple modalities (e.g., vision, audition, and somatosensation) and translate sensory signals into motor commands used to guide orienting movements. Primary outputs from the superficial region include projections to the pulvinar and lateral intermediate areas of the thalamus, which in turn project to areas of the cerebral cortex that are involved in controlling eye movements. The projections from the deeper layers have also been characterized extensively, which include two descending projections, traveling to the brainstem and spinal cord, and ascending projections to multiple sensory and motor centers (Benavidez et al., 2021). Given its capacity to register multiple sensory modalities and to coordinate signals into appropriate motor commands, the SC has been widely studied for investigating the principles underlying both multisensory and sensorimotor processing.

During nervous system development, axon guidance and targeting selection are critical steps toward establishing proper neural circuits. The SC has been a prominent model for investigating the mechanisms by which axons innervate their target regions. Accordingly, studies have revealed that within each SC lamina, RGC inputs are mapped topographically with respect to the visual field; the temporal–nasal axis of the retina projects along the anterior–posterior axis of the SC, and the dorsal–ventral axis of the retina projects along the lateral–medial axis of the SC. The formation of topography along these axes is a result of combined effects of molecular labels, such as Eph receptors and ephrin (Cang et al., 2008; Feldheim and O’Leary, 2010; Huberman et al., 2008a; Rashid et al., 2005). Others have used transgenic mice to demonstrate that different RGC types project to distinct SC regions and provide inputs to multiple types of SC neurons (Huberman et al., 2008b; Kim et al., 2010; Martersteck et al., 2017).

Single cell RNA sequencing (scRNAseq) and single nucleus RNA sequencing (snRNAseq) have been widely used in the central nervous system (CNS) to characterize cellular heterogeneity and the dynamic changes of specific cell populations over time. To define distinct cell types and cellular subtypes of the SC at the genome-wide level, and their changes across development, here we performed snRNAseq analysis of the mouse SC isolated at an embryonic stage and at various ages during the first month of postnatal life. Our results revealed molecularly distinct excitatory and inhibitory neuronal subtypes, many of which reside exclusively in specific SC laminae or project to specific brain region. Moreover, we find numerous genes that are differentially expressed in neurons and glial cells across the different postnatal ages, many of which are known to regulate axon guidance, targeting, and synapse formation. For convenient access to these data, we have created a user friendly web portal that provides a platform for analyzing and comparing gene expression profiles in the SC cell types and neuronal subtypes across the ages: https://parklabmiami.shinyapps.io/superior-colliculus-snRNAseq/

## RESULTS

### Single-Nucleus Profiling of Developing and Mature Mouse Superior Colliculus

Some cell types, such as neurons, are more susceptible to tissue dissociation processes and are underrepresented in scRNA-seq data. In contrast to whole cells, nuclei are more resistant to mechanical assaults and can be isolated from frozen tissue (Bakken et al., 2018). Indeed, studies have revealed that snRNA-seq profiling can provide an unbiased survey of neural cell types and as a result, snRNAseq has become a popular choice for profiling susceptible cells such as neurons (Velmeshev et al., 2019).

To profile gene expression of SC during development and maturation, we analyzed the transcriptomes of single nuclei across four time points: embryonic day 19 (E19), postnatal day (P) 4, P8 and P21. We selected these time points because together they encompass discrete yet overlapping developmental events including axonal outgrowth, axonal targeting, topographic mapping, synaptogenesis, oligodendrocyte differentiation, myelination, and synaptic refinement and maturation. For each age, we micro-dissected the SC territory from several animals, which were pooled. After detergent-based digestion and mechanical dissociation followed by nuclei isolation using a sucrose density gradient centrifugation, the single nucleus suspension was processed and sequenced using the 10X Genomics droplet-based snRNA-seq platform (Figure 1A). Downstream clustering and gene expression analyses were done using Seurat and the Bioconductor suite of bioinformatics tools. After stringent quality control (Figure S1), a total of 9,728 high-quality nuclei (2,585 from E19, 2,386 from P4, 2,508 from P8, and 2,249 from P21) were retained for downstream analysis. An average of 10,451 unique molecule identifiers (UMIs) were captured per nucleus (7,457 for E19, 9,882 for P4, 10,961 for P8, and 13,928 for P21) and an average of 3,535 genes were detected per nucleus (3,030 for E19, 3,574 for P4, 3,644 for P8, and 3,953 for P21) (Figure S1B).

**Figure 1.**
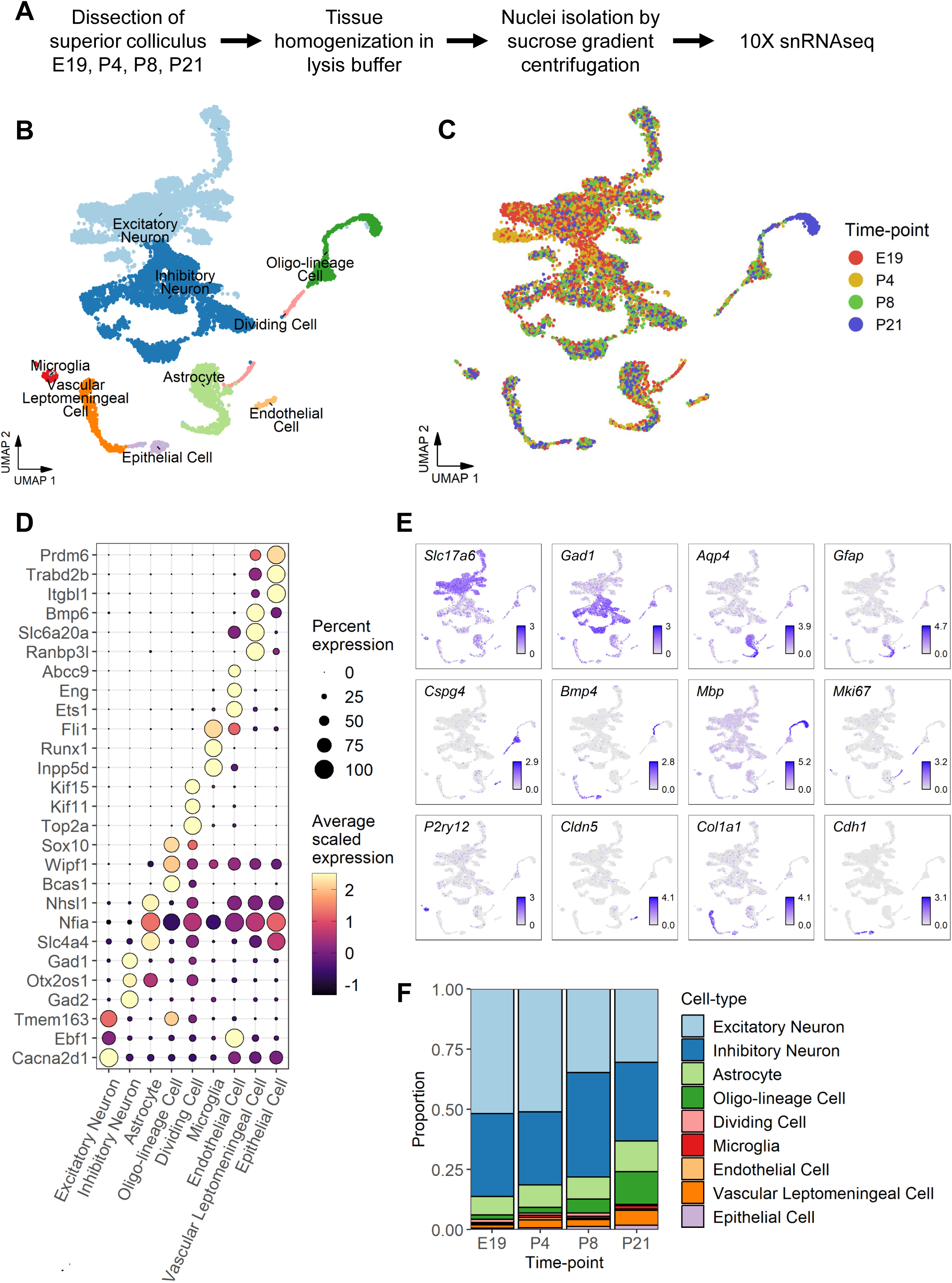
Single nucleus RNA-sequencing of developing superior colliculus. **A**) Overview of experimental design. **B**) Summary Uniform Manifold Approximation Projection (UMAP) plot of all major cell types identified from integrated analysis. Dots represent gene expression profiles for individual nuclei and are colored by cell type. **C**) UMAP of nuclei colored by developmental time point of origin. **D**) Dot plot of top differentially expressed genes per cell type. **E**) Marker genes identified previously from the literature used to annotate cell types. **F**) Quantification of cell type proportions across different ages.

### Identification of Cell Types in the Developing Mouse SC

To identify shared and unique cell types across SC development, we used the Seurat integration pipeline to jointly analyze cells from all four time points and perform unsupervised clustering (see Methods). Projection of cells onto UMAP plots (Becht et al., 2018) revealed structures suggestive of cell lineage and differentiation (Figures 1B and 1C). Using the top differentially expressed genes (DEGs) per cluster and a panel of previously described cell type marker genes (e.g. *Slc17a6*, *Gad1*, *Aqp4*, *Gfap*, *Cspg4*, *Bmp4*, *Mbp*, *Mki67*, *P2ry12*, *Cldn5*, *Col1a1*, and *Cdh1*; Figures 1D and 1E), we identified a total of 31 cell clusters which were divided into 21 neuronal and 10 non-neuronal clusters. These clusters represented most major cell types that are known to comprise the SC including neurons, astrocytes, microglial cells, and oligodendrocyte-lineage cells, as well as endothelial and vascular leptomeningeal cells (Figures 1B and 1D).

Consistent with the notion that neuronal nuclei are expected to survive the dissociation protocol, neurons made up a sizable portion of the cells, constituting about 77% of all cells identified. Among all neuronal cells, 54% were excitatory neurons and 46% were inhibitory neurons but this ratio varied by time point. While neurons at E19 and P4 were approximately 60% excitatory and 40% inhibitory, neurons at P8 and P21 were approximately 46% excitatory and 54% inhibitory (Figure 1B and 1F). Consistent with the timing of oligodendrocyte differentiation and myelination, the proportion of *Mbp*^+^ oligodendrocyte-lineage cells, or mature oligodendrocytes, were higher in P8 and P21 than E19 and P4 (Figures 1C and 1F). We also observed that the proportion of non-neuronal cells, such as microglia and astrocytes, gradually increased over time (Figure 1F). The temporal trends we observed in the data agreed with many previous studies on brain cell type composition and development, speaking to the quality of the cells captured (Kalish et al., 2018; Velmeshev et al., 2019; Xie et al., 2021). With these recapitulated trends affirming data quality, we next proceeded to further analyze individual cell types through separate analyses.

### Heterogeneity of SC neurons across development

Several studies have described molecularly distinct types of SC neurons, but a genome-wide reference of neuronal subtypes throughout SC development has been lacking. Because neurons made up a sizable percentage of the sequenced nuclei in our dataset, we were well positioned to provide a reference map of the transcriptional heterogeneity among SC neuronal subtypes. To this end, we performed a nested cluster analysis using only neurons so that only genes variably expressed amongst neurons were considered. This resulted in 25 neuronal clusters (Figures 2A and S2A) which, when projected onto a UMAP (Figure 2A), were principally segregated by excitatory or inhibitory subtype as confirmed by expression of the gene *Slc17a6*, which encodes the vesicular-glutamate transporter VGLUT2, or by expression of glutamate decarboxylase genes, *Gad1* and *Gad2*, respectively (Figures 2C and 2D). These neuron clusters were therefore annotated as EN1-13 or IN1-12 subtypes. We then pivoted the data to measure neuronal subtype quantities across developmental ages and found that most subtypes were detected at most time points (Figures 2B, S2B and S2C). However, subtypes EN1 and IN1 were preferentially enriched at earlier time points E19 and P4, and many subtypes increased in proportion at later P8 and P21. At the extreme, some subtypes were temporally restricted, such as subtype EN7, which was found only at E19 and P4 stages (Figure S2C).

**Figure 2.**
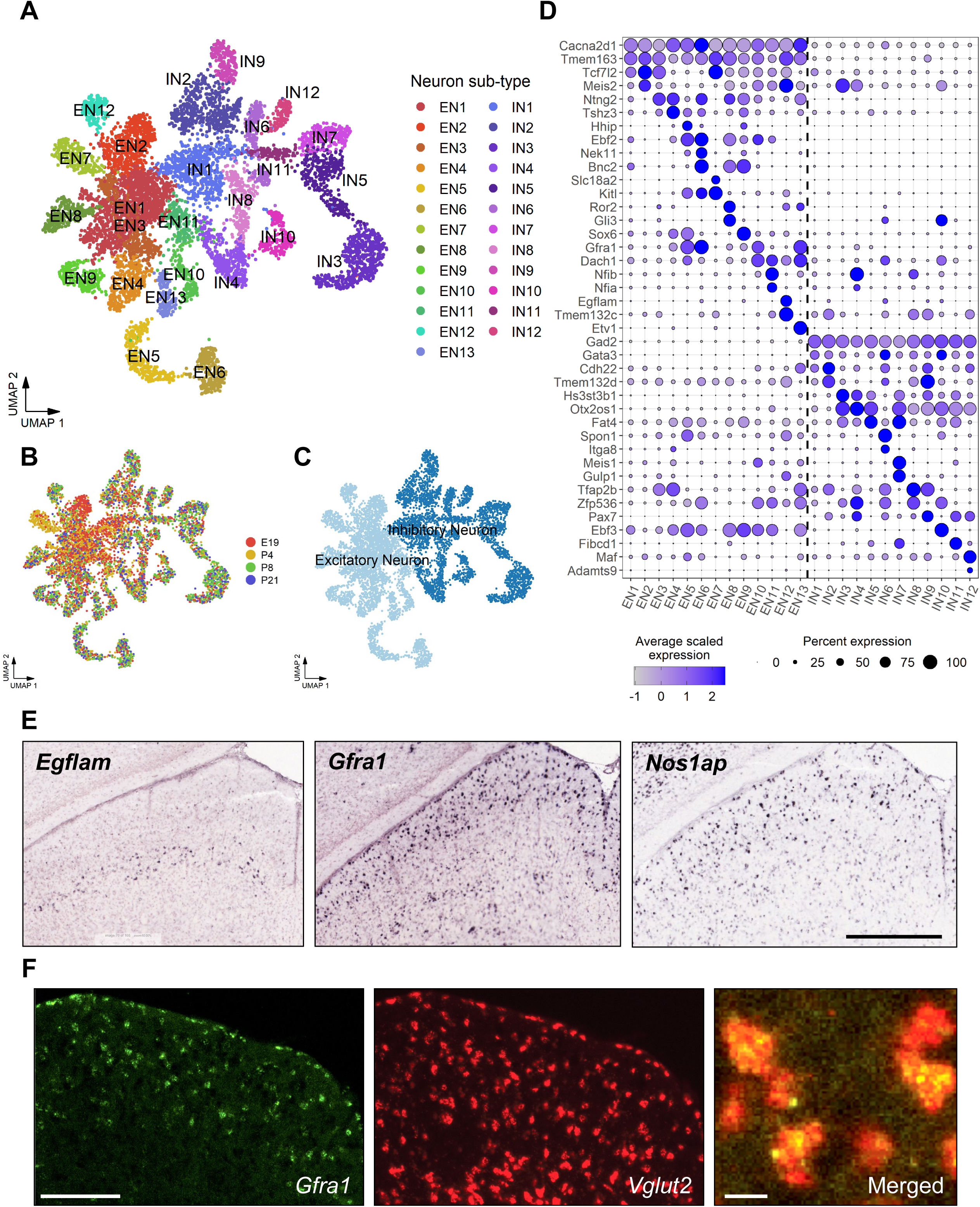
Cluster analysis of neurons reveals transcriptionally and temporally defined neuronal subtypes. **A**) UMAP of neuronal subtypes identified through cluster analysis of neuronal cells. Cells are colored by subtype. EN = Excitatory Neuron subtype. IN = Inhibitory Neuron subtype. **B**) UMAP of neurons colored by sample developmental time point. **C**) UMAP of neurons colored by principal neuron class. **D**) Dot plot of the top differentially expressed genes for each neuronal subtype compared against all other neurons combined. Dashed line partitions excitatory and inhibitory neuronal subtypes. **E**) In situ hybridization against *Egflam* (left), *Gfra1* (center), and *Nos1ap* (right) from the Allen Brain Atlas. Genes were identified from the list of subtype differentially expressed genes as shown in Figure 2B. **F**) Representative coronal mouse brain sections showing SC. Fluorescent in situ hybridization was performed using probes against *Gfra1* (green) and *Vglut2* (red). Scale bars, 500 µm in E and F (left two panels), and 10 µm (right panel).

Subsequent differential expression analysis revealed both distinct and overlapping marker genes across subtypes (Figure 2D). While some neuron subtypes could be uniquely identified by the expression of a single gene, such as *Etv1* expression by EN13 and *Maf* by IN12, other subtypes were better described using combinations of multiple genes, similar to the approach taken in previous studies that classified SC neurons using multiple molecular markers (Byun et al., 2016). For example, subtype EN7 can be specified by joint expression of *Kitl* and *Slc18a6,* and subtype IN10 can be specified by joint expression of *Ebf3* and *Gata3.* Given the varying degree of marker specificity, we next quantified subtype expression profile similarities through a dendrogram analysis (Figure S2D). While our results re-emphasized principal neuronal class (excitatory vs inhibitory) as the greatest source of variation, we also observed that some neuron subtypes, such as EN1 and IN1, were less distinct than others. We confirmed this observation by counting the number of DEGs per neuronal subtype grouped by neuronal class and found that, indeed, subtypes EN1 and IN1 had the fewest numbers of DEGs while other, more distinct subtypes had upwards of 100 DEGs (Figures S2E and S2F). Combined with our proportions data, we speculate that subtypes EN1 and IN1 represent less differentiated neuron subtypes at earlier time points that become more differentiated subtypes throughout development.

### Correlates between excitatory subtype markers and SC laminae

Previous studies have demonstrated that morphologically and physiologically distinct types of excitatory SC neurons are localized in distinct laminae and contribute to various behavioral responses in mice (Ito and Feldheim, 2018; Wheatcroft et al., 2022). From our initial differential expression analysis, we next performed a more detailed inspection of the marker genes that distinguish the excitatory subtypes. Among the 13 excitatory neuronal subtypes, we found subtypes that exhibit enriched expression of genes that are already known to label subpopulations of SC neurons (Figure 2D). Subtype EN13 expressed *Etv1* exclusively, a gene known to be expressed in a subpopulation of neurons in the SO layer (Byun et al., 2016). *Gfra1* was also highly expressed in this subtype. A previous study has shown that *Pitx2* defines a subpopulation of excitatory neurons in the stratum griseum intermediate (SGI) (Masullo et al., 2019). We found that subtype EN8 expressed elevated levels of *Pitx2* as well as *Gli3* and *Rorβ* compared to other excitatory subtypes (Figures 2D and S6E). Thus, EN8 likely represents this previously characterized SGI cell type. Other subtypes with notable gene enrichment were EN12, which was enriched for *Egflam and Tmem132c* expression, and EN11 which was enriched for *Nfib* and *Nfix*.

Given the topographic arrangement of SC neurons, we next examined whether these neuronal subtypes were localized in discrete laminae in adult SC. In the Allen Brain *in situ* hybridization database, *Gfra1,* which we found to be expressed in excitatory neurons and particularly high in EN6, was detected primarily in the SGS (Figures 2E and 2F). *Egflam,* a marker for EN12, was detected predominantly in the SO (Figure 2E). In the deeper layers, *Pde1c*, high in EN7, was detected predominantly in the SGI (data not shown). To validate the expression of these marker genes in the mouse tissues, we performed fluorescent in situ hybridization (FISH) against *Gfra1*. Consistent with our snRNAseq data, *Gfra1* was detected predominantly in the SGS, and these *Gfra1*^+^ cells expressed the excitatory marker *Vglut2* (Figure 2F).

### Correlates between inhibitory neuron subtype markers and SC laminae

Just as in the clusters of excitatory neurons, our differential expression analysis identified 12 distinct inhibitory neuronal subtypes with characteristic transcriptional signatures (Figure 2D). Notably, *Spon1* was specific to IN6, and *Meis1* and *Gulp1* were specific to IN7. Subtype IN11 could be defined by high expression of *Fibcd1* and *Pax7.* We examined whether these subtypes reside in distinct layers in the SC. As in the excitatory neurons, markers of several inhibitory neuronal subtypes showed layer specific localization. For example, we detected *Nos1ap,* a marker for IN3, primarily in the SGS (Figure 2E). Together, our results describe the genetic heterogeneity in neurons during SC development and illustrate that neurons with distinctive transcriptomic profiles are often located in different layers of the SC.

### Changes in neuronal gene expression across development

Expression of various genes including transcription factors and cell adhesion molecules, is developmentally regulated. Protein products of these genes are known to regulate critical neurodevelopmental processes including axon growth, axon targeting, retinotopic mapping, and synapse formation and maturation. Thus, we sought to identify genes that were differentially expressed across the four time points in our data. Given the distinction between excitatory and inhibitory neurons, we first evaluated for temporal changes within each class individually as in our cluster analysis.

For excitatory neurons at E19, the top enriched genes included *Nnat, Nsg1, Nsg2, Fam171a2, Basp1, Pcsk1n,* and *Rtn1* (Figure 3A). At P4, transcription factors *Zic1* and *Tcf7l2*, whose gene product plays a key role in Wnt signaling, were highly upregulated. Notably, inspection of Allen Brain *in situ* data showed that, at E18.5 and P4, *Zic1* expression was high only within the medial side of the SC in the SGI layer, and not present in the lateral side (Figure 3D, top 3 panels). By adulthood (i.e. P56), medial specific *Zic1* expression in the SC was no longer detected (Figure 3D, bottom panel). This expression pattern suggests that *Zic1* may determine axon directionality along the lateral-medial axis in this region (see Discussion). At P8, the top upregulated genes included transcription factor *Lef1,* and guidance molecules *Cdh13* and *Dscaml1*. Interestingly, within the midbrain, *Lef1* expression was almost exclusive to the SC, suggesting that Lef1 may have specific functions for the SC neurons (Figure 3E). Notably, we observed that the greatest number of gene expression changes occurred between P8 and P21 (Figure 3C), where P21 was enriched for genes such as *Lncpint, Specc1, Rasgrf1, Naaladl2, Aifm3,* and *Pcgf5*.

**Figure 3.**
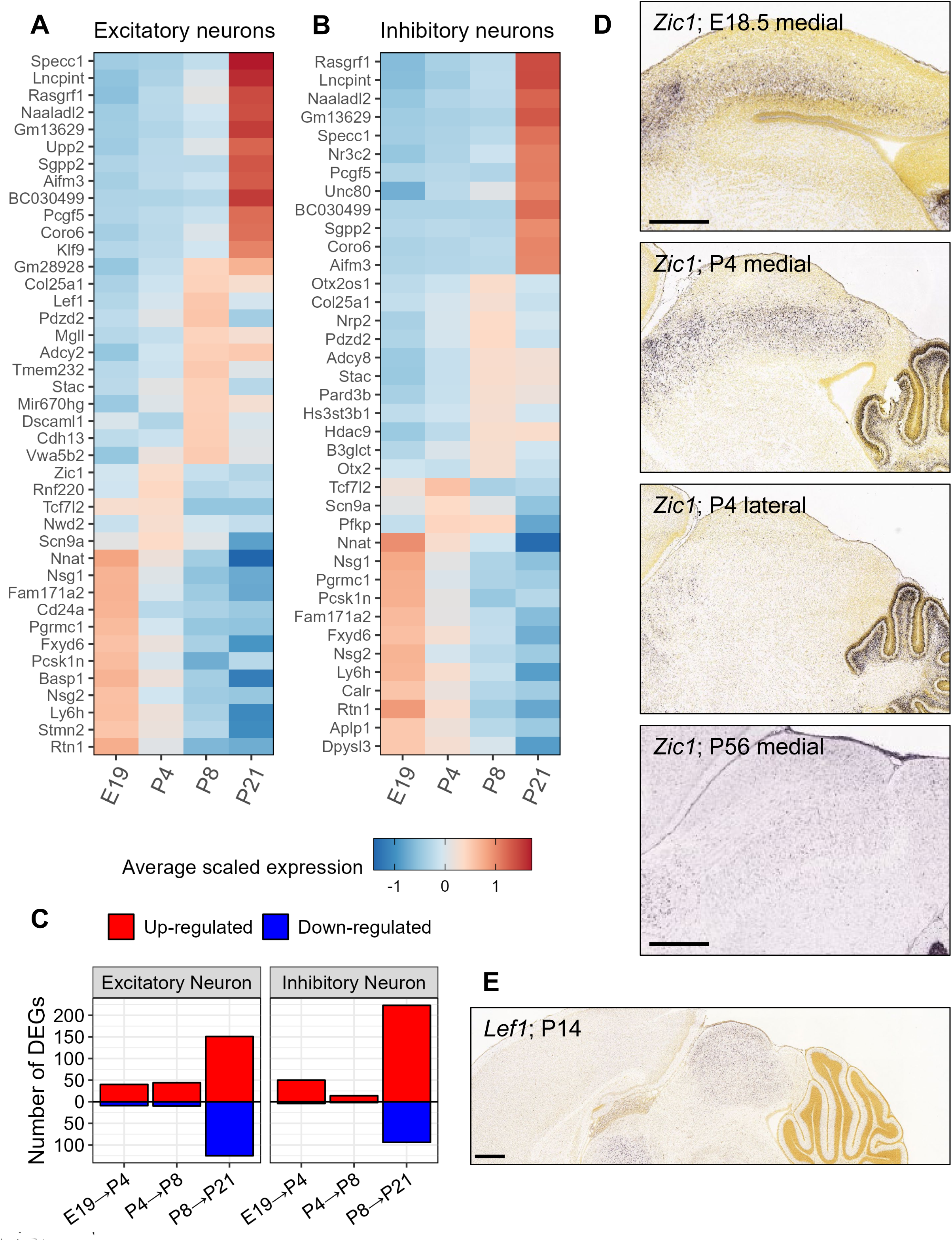
Identification of transcriptional changes across neurons across development. **A** and **B**) Heatmap of the top differentially expressed genes per developmental time point in **A**) excitatory neurons or **B**) inhibitory neurons. Excitatory or inhibitory neuron subtypes were aggregated prior to differential expression testing. **C**) Quantification of the number of up-regulated or down-regulated differentially expressed genes between sequential developmental time points. **D**) In situ hybridization against *Zic1* from Allen Brain Atlas. **E**) In situ hybridization against *Lef1* from Allen Brain Atlas. Wilcoxon Rank Sum test as implemented in Seurat’s `FindMarkers()`function. Scale bars, Scale bars, 500 µm.

Within inhibitory neurons, we performed a similar series of differential expression tests comparing enriched genes at each time point (Figure 3B). We found a large degree of similarity in temporal expression patterns between inhibitory and excitatory neurons. For example, as is the case for excitatory neurons, genes such as *Nnat, Nsg1, Nsg2, Pcsk1n,* and *Rtn1* were enriched at E19, and the transcription factor *Tcf7l2* was enriched at P4. Furthermore, we observed that the largest number of gene expression changes took place between P8 and P21. We did, however, also identify DEGs that were unique to inhibitory or excitatory neuron development. Whereas genes such as *Gfra1* were upregulated at P8 in excitatory neurons, genes such as *Otx2* and *Otx2os1* were upregulated at P8 compared to other time points and were unique to inhibitory neurons. It is worth noting that these genes corresponded to markers of neuronal subtypes whose proportions varied over time (Figure S2C).

Given this high overlap in temporal patterns, we next combined excitatory and inhibitory neurons as a single group to identify temporal patterns that were globally shared among all neurons (Figure S3A). Our analysis confirmed our previous findings; we found *Basp1*, which encodes a protein known for its role in neurite outgrowth (Korshunova et al., 2008), to be enriched at E19. We again identified the key Wnt pathway transcription factor, *Tcf7l2*, to be enriched at P4. Accordingly, further characterization through Gene Ontology (GO) analysis demonstrated down-regulation of genes associated with Wnt signaling from P4 to P8, supporting the importance of Wnt signaling during this period (Figure S3B). Moreover, these neurons were most transcriptionally dynamic between P8 and P21, and upregulated genes outnumbered downregulated genes in each adjacent age comparison (Figure 3C). The overt changes during this period may reflect neuronal developmental processes associated with synaptic maturation and the appearance of oligodendrocytes and myelination (Figure 1F). Indeed, our GO analysis of genes upregulated from P8 to P21 revealed terms such as “regulation of synaptic plasticity” and “synaptic vesicle exocytosis” (Figure S3B). Overall, both excitatory and inhibitory neurons showed similar temporal patterns of transcriptional activity whose inferred biological functions through GO analysis recapitulate known developmental milestones, lending further validity to our dataset.

### Changes in glial gene expression across development

In addition to neurons, we successfully captured virtually all glial cells, of which astrocytes were the most abundant. Comparisons of astrocyte gene expression across the developmental time points (Figure S4A) revealed an enrichment of axon guidance molecule genes, such as *Ephb1*, *Robo1,* and *Ephb2*, at E19 and their decreased expression over time. Additionally, P4 astrocytes highly expressed *Hs6st3*, a heparan sulfate (HS) sulfotransferase known to modify HS and regulate adhesion. P8 astrocytes were enriched for *Ptchd4*, a repressor of canonical hedgehog signaling, and glutamate metabotropic receptor *Grm7*. Like neurons, astrocytes also demonstrated a large increase in the number of DEGs between P8 and P21 (Figure S4B). Among them were genes such as *Gja1* and *Gjb6*, encoding the gap-junction proteins also known as Connexin-30 and Connexin-43, respectively, which map to the GO term “cell junction assembly” (data not shown). These genes have been implicated in astroglial synapse coverage (Pannasch et al., 2014). Other genes important for astrocytic function, such as those that encode ion channels and ion channel regulators (e.g. *Trpm3 and Lgi1*) and phospholipases (e.g. *Plcb1 and Gpld1*), were also upregulated at P21. These results collectively suggest that astrocytes may play a role in axon guidance earlier in development and reaffirm their roles in synaptic refinement in young adulthood concurrent with the appearance of mature *Mbp*^+^ oligodendrocyte-lineage cells by P21.

Oligodendrocyte-lineage cells showed UMAP coordinates that were suggestive of differentiation across development (Figure 1B). In agreement with this observation, our differential expression analysis showed an enrichment of oligodendrocyte progenitor cell (OPC) marker genes such as *Olig1* and *Pdgfra* at E19 and a strong upregulation of genes whose products encode for components of the myelin sheath such as *Mag*, *Plp1*, *Mbp*, and *Mog* at P21 (Figure S4C). Moreover, we observed that *Sox6* expression remains high from E19 to P8 but is downregulated at P21. This is consistent with the notion that Sox6 maintains the precursor state of oligodendroglial cells, thereby ensuring the proper timing of myelination in the CNS (Ittner et al., 2021). Other genes showed a similar expression pattern, such as *Lrrtm4* and phosphodiesterase *Pde4d*. Overall, like for other cell types, we observed substantially more genes differentially expressed between P8 and P21 than compared to other time points, again suggesting that this critical period is highly transcriptionally dynamic (Figure S4D).

Though we successfully identified a microglia cluster, they totaled 90 nuclei, reducing our power to identify DEGs. Thus, their analysis was not included in this report.

### Comprehensive assessment of major axon guidance molecules

#### Cadherins and protocadherins

Cadherins play important roles in axon guidance and target selection. Among the cadherin family members, we found that the expression of *Cdh8, Cdh12, Cdh13,* and *Cdh18* was particularly high in neurons and low in non-neuronal cells (Figure 4A). In contrast, we observed significant expression of specific cadherin genes in non-neuronal cells; *Cdh20* expression was exceptionally high in astrocytes, whereas *Cdh1* expression was specific to epithelial cells. During our analysis, we noticed that while most cadherins and protocadherins were expressed at similar levels in excitatory and inhibitory neurons as a class, their expression varied highly at the subtype level. Since selective cadherin expression has been shown to promote neuron class-specific cell adhesion and synaptic connection (Jontes, 2018; Sanes and Zipursky, 2020), we generated a comprehensive panel of cadherin family gene expression across neuronal subtypes. From this panel, we found that *Cdh7* was expressed at higher levels in subsets of excitatory neurons (EN5 and EN6) (Figure 4B). Previously, Byun et al. described *Cdh7* as a marker for a subpopulation of neurons in the upper SGS (uSGS) (Byun et al., 2016). Our data extend their findings and demonstrated that *Cdh7* marks a subpopulation of excitatory neurons in the uSGS. Conversely, we found that *Cdh12* and *Cdh22* were preferentially enriched in inhibitory subtypes compared to excitatory populations.

**Figure 4.**
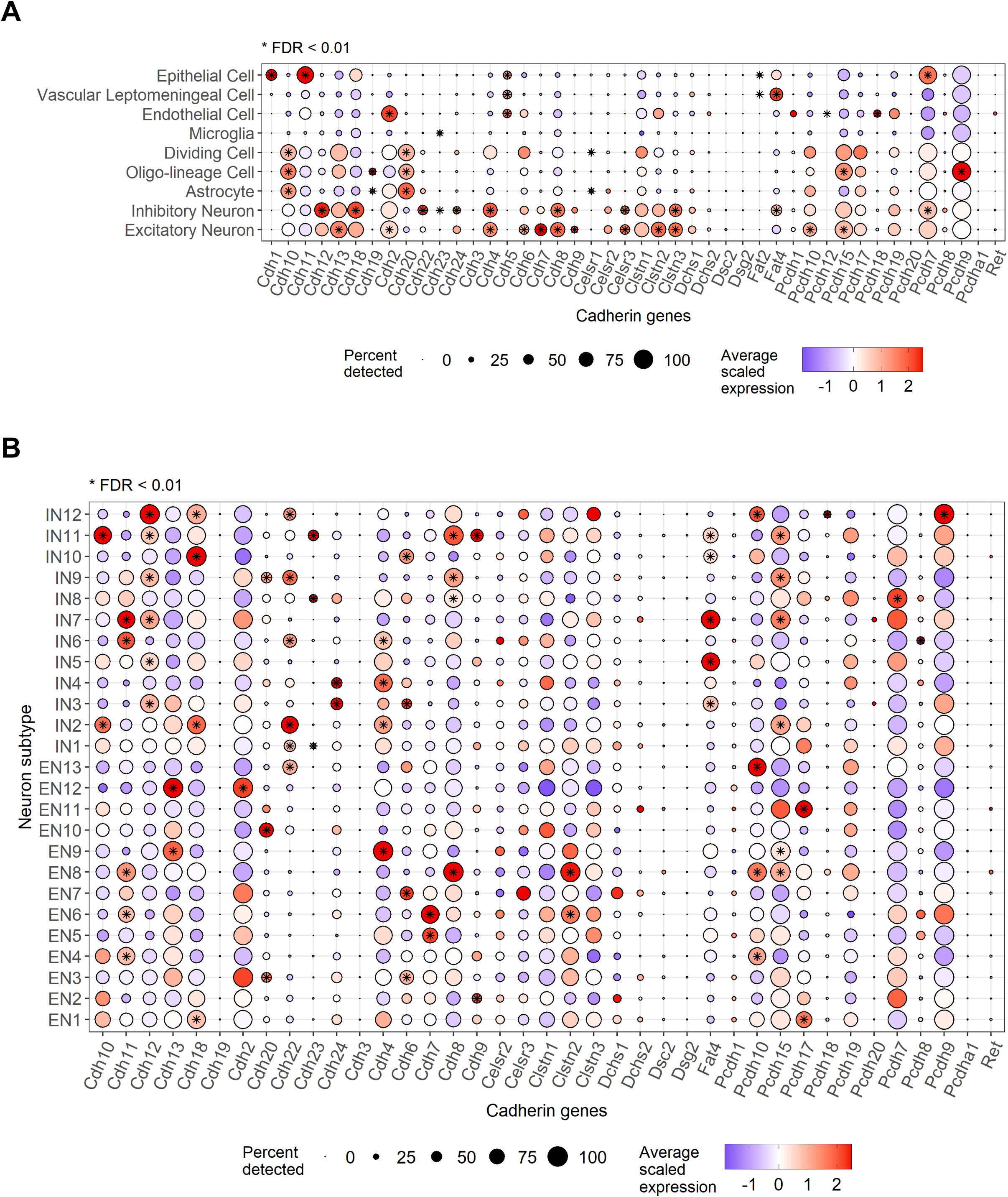
Comprehensive panel of cadherin family gene expression across superior colliculus cell types and neuronal subtypes. **A**) Dot plot of scaled gene expression values of cadherin family genes across SC cell types. Only genes which were detected in at least 0.1% of all cells are shown. **B**) Dot plot of scaled gene expression values of cadherin family genes across SC neuron subtypes. Only genes which were detected in at least 0.1% of all neuronal cells are shown. Stars indicate statistically significant enrichment of each cadherin gene in each group compared to all other cells combined. Wilcoxon Rank Sum test as implemented in Seurat’s `FindMarkers()` function. FDR = false discovery rate.

#### Eph receptors and ephrins

This family of receptors and ligands are key signaling molecules involved in retinocollicular topographic mapping. Accordingly, many Eph receptors were highly detected in the SC neurons across multiple time points with the exception of *Epha1, Epha2, Ephb3* and *Ephb4* (Figure S5A). Multiple ephrin ligands were also detected, and some, such as *Efna2* and *Efnb3,* showed downregulation in neurons over time. Overall, we observed that genes from the Eph receptor and ephrin family were primarily expressed in neurons compared to non-neuronal cells. Some exceptions to this included astrocytic expression of *Epha5, Ephb1,* and *Ephb2*, which decreased significantly over time, and *Efnb3* expression by oligodendrocyte-lineage cells.

#### Semaphorins and plexins

These molecules regulate functions related to cell morphology and communication that contribute to axon guidance. In our data, most semaphorin-plexin family genes were detected in SC neurons across all time points such as *Plxna4*, *Sema4f*, and *Sema4g*. (Figure S5A). Whereas neurons maintained constant expression levels of most of these genes, *Sema3f* and *Plxna1* showed a significant decrease across developmental time points. However, non-neuronal cells also expressed members of the semaphorin-plexin family. For example, astrocytes demonstrated enriched expression of *Plxnb1* and a strong upregulation of *Sema4b* over time, and oligodendrocyte-lineage cells showed similarly strong temporal changes in *Plxnb3*, *Sema4d*, and *Sema6d*.

#### Robo receptors and Slit

While transcripts for *Robo1* and *Robo2* were detected in nearly all cell populations, which we speculate is due to ambient RNA contamination (see Methods), these genes were most highly expressed in SC neurons. In contrast, *Robo3* was detected at low levels in all cell types at all time points except for in excitatory neurons, whose expression significantly increased across the time points. Further inspection of *Robo3* showed that its expression was limited to neuron subtypes EN5 and EN6, subtypes whose proportions correspondingly increased over time. These same *Robo3*^+^ subtypes comprised a subset of *Gfra1*^+^ neurons which we have shown to be enriched in the SGS (data not shown). Finally, while the Robo ligands *Slit2* and *Slit3* were most highly detected in epithelial cells and only moderate expressed by SC neurons, *Slit1* was almost exclusively expressed by neurons. (Figure S5A).

### Correspondence between neuronal subtypes and published SC scRNAseq data

Several recent studies have profiled murine SC neurons using scRNAseq or snRNAseq. Two of these studies elegantly utilized transsynaptic and retrograde AAVs to specifically profile retinorecipient neurons and SC projection neurons, respectively. Tsai et al. developed the Trans-seq method which used modified adeno-associated virus (AAV)-wheat germ agglutinin (WGA) to identify and profile SC neurons that are post-synaptic to RGCs (Tsai et al., 2022). Cheung et al. developed Vector-seq using retrograde viruses to profile SC projection neurons that send axons to different brain regions (Cheung et al., 2021). In both studies, the authors provided user-defined subtypes of SC neurons, but how these subtypes compare across studies is not known, and whether these populations are detected in our development time-course dataset is unclear.

To elucidate unified molecular definitions of SC neuron subtypes, we pulled data from each study and first asked whether neuronal subtypes as defined in our study were detected in these published data. To this end, we applied the `SingleR` classification algorithm (Aran et al., 2019) (Figure 5A) using the snRNAseq data in our study as a reference and the Trans-seq and Vector-seq data as query sets. Briefly, the `SingleR` algorithm computes Spearman correlations between cells of the query data and cells of the reference data on the subset of marker genes for subtypes of the reference data i.e., EN1-13 and IN1-12 subtype marker genes. Per-subtype scores are determined from these correlations, and the subtype with the highest score becomes the predicted subtype (see Methods for further details). This approach is especially suitable for this analysis given the technical noise driven by ambient RNA contamination (see Methods), differences in sequencing depth and dissociation method, and study-associated batch effects.

**Figure 5.**
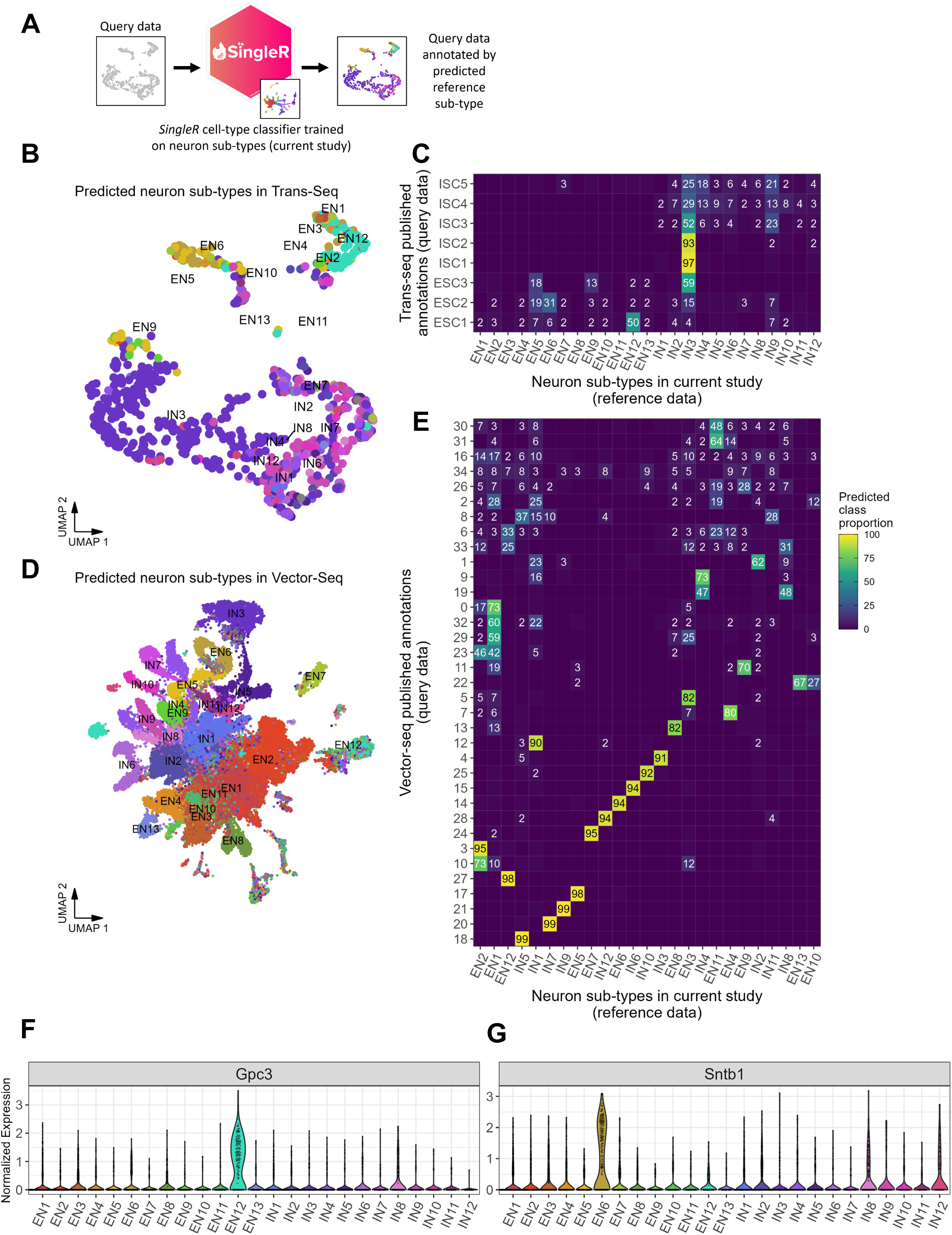
Comparison of neuron subtypes against published superior colliculus sn-or scRNAseq data. **A**) Schematic of `SingleR` approach to classify neurons from published data sources using marker genes of neuron subtypes defined in our analysis. Briefly, for each query cell, *SingleR* computes correlations between its expression profile and the average expression profile for each reference label (i.e. our sub-types). Query cells in the upper quantiles of per-label correlations are annotated as that label. **B**) UMAP of cells from Trans-Seq data (see Figure 5B by Tsai *et al*., 2022) or **D**) Vector-Seq data (see Figure 3B by Cheung *et al*., 2021) colored by predicted classification. **C**) Heatmap of correspondence between neuron sub-types identified in current analysis and clusters identified by Tsai *et al*. in Trans-Seq, or **E**) clusters identified by Cheung *et al*. in Vector-Seq. Heatmap values depict the proportion of cells from Y-axis clusters that were annotated as X-axis neuron subtypes *e.g.* 99% of cells in cluster 18 from Vector-Seq were classified as IN5 based on marker genes for IN5. Sums along rows equal to 100. **F** and **G**) Violin plot of *Gpc3* **F**) and Sntb1 **G**) expression in excitatory and inhibitory neuron subtype. Clusters Ex-6 (i.e. Gpc3+ neurons) and Ex-8 (i.e. Sntb+ neurons) from Xie et al. correspond to our EN12 and EN6, respectively.

The original Trans-seq study categorized retinorecipient SC neurons into three excitatory subtypes (ESC1-3), and five inhibitory subtypes (ISC1-5) (Tsai et al., 2022). To compare these subtypes against subtypes from our study, we applied `SingleR` and computed the proportions of predicted labels that comprised each of the Trans-seq subtypes (Figures 5B and 5C). Our analysis showed that more than 93% and 97% of cells in Trans-seq ISC1 and ISC2 subtypes, respectively, were predicted to be our inhibitory subtype IN3. This suggests a high degree of similarity between these three populations and perhaps that IN3 may be a retinorecipient population. Conversely, only excitatory neuron subtypes EN5, EN6, EN9, and EN12 had predictions that comprised more than 10% of any Trans-seq subtype. This is in line with the expectation that retinorecipient neurons comprise only a subset of all SC neurons and further suggests that RGCs only project onto SC neurons in a subtype-specific manner. Overall, five neuronal subtypes (IN3, IN9, EN12, IN4, and EN5) comprised approximately 75% of predicted subtypes in the Trans-seq dataset (Figure S6A), indicating that these subtypes may represent most likely candidates of RGC post-synaptic input.

The Vector-seq study by Cheung *et al*. used snRNAseq to profile about 55,000 nuclei of the SC. When we applied the same `SingleR` approach to these data, we found strong concordance between our neuron subtypes and Vector-seq clusters. Thirteen of 35 Vector-seq clusters had greater than 90% of its cells predicted to be of a single neuron subtype as defined in our study and 20 of 35 clusters were comprised of more than 70% of a single subtype (Figures 5D and 5E). Notably, using their retrograde labeling strategy, Cheung *et al*. reported that approximately 1,500 Vector-seq excitatory neurons (clusters 2 and 7, Cheung *et al*. figure 3C) were SC neurons that project to contralateral paramedian pontine reticular formation (PPRF) (i.e. SC neurons that are known to control orientating movements), and about 3,100 cells were SC neurons (clusters 10 and 11, Cheung *et al*. Figure 3C) that project to the thalamic lateral posterior nucleus (LP). We repeated `SingleR` analysis on only excitatory neurons and found that Vector-seq excitatory neuron cluster 7 had high concordance with our excitatory subtype *Pitx2*^+^ EN8, suggesting that EN8 likely represents SC neurons projecting to the PPRF (Figures S6B and S6E). Moreover, Vector-seq excitatory neuron cluster 10 (which is *Ntng2*^+^) was comprised mostly of *Ntng2*^+^ excitatory subtype EN3, while Vector-seq excitatory neuron cluster 11 was made up of a combination of EN2 and EN12, which could be distinguished by *Cbln2* (Figures S6B and S6F). From this, we speculate that subtypes EN3, EN2, and EN12 are SC neurons which project to the thalamic LP.

Together, these results highlight the importance of a unified, reference set of SC neurons that can extend findings from viral tracing studies to infer involvement of specific neuronal subtypes in neural circuitry.

Using in situ hybridization and immunostaining, Byung *et al*., demonstrated that several molecular markers collectively categorize 10 superficial SC (sSC) neuronal types. These genes include *Etv1*, and *Rorβ.* As aforementioned, we found that *Etv1* expression was unique to EN13. Several subtypes from both excitatory and inhibitory populations in our study expressed *Rorβ*, albeit the expression was more widespread in the excitatory neuron clusters (data not shown). Xie *et al*. performed snRNAseq on SC from adult mouse brains and clustered the excitatory and inhibitory neurons into 9 and 10 subtypes, respectively (Xie et al., 2021). We observed good correspondence among many neuronal subtypes between the two studies, demonstrating the validity of these snRNAseq data. For example, clusters Ex-6 (i.e. *Gpc3*^+^ neurons) and Ex-8 (i.e. *Sntb^+^* neurons)(Xie et al., 2021)from Xie *et al*. corresponded to our EN12 and EN6, respectively (Figures 5F and 5G).

Previous studies have also classified inhibitory SC cells into three non-overlapping categories in mice; those that express *Sst, Vip,* or *Pvalb.* As for the inhibitory neurons identified in this study, IN4, IN8, and IN12 corresponded to *Sst^+^* neurons, and IN9 corresponded to *Vip*^+^ neurons. *Pvalb*^+^ neurons, which are found in the lower SGS and SO (Illing et al., 1990), innervate the lateral posterior nucleus of the thalamus. Thus, some *Pvalb*^+^ neurons are projection neurons, as opposed to interneurons (Casagrande, 1994; Mize, 1996). While *Pvalb* marks a specific subclass of GABAergic fast-spiking inhibitory interneurons in the cerebral cortex, striatum, and hippocampus (Hu et al., 2014), it was shown that *Pvalb*^+^ neurons in the sSC present heterogeneous spiking profiles and morphologies, and only a fraction contained GABA. On the other hand, a higher proportion of the *Pvalb*^+^ population in the intermediate layers of SC showed the presence of GABA. Consistent with the notion that *Pvalb*^+^ neurons in the SC are likely heterogeneous populations of both glutamatergic and GABAergic neurons (Villalobos et al., 2018), we observed *Pvalb* expression in various neuronal subpopulations in our data (data not shown).

### Characterization of a subpopulation of GABAergic neurons: Pax7-expressing neurons

The SGS contains a high density of GABAergic neurons; 30% of all neurons in this region are GABAergic (Mize et al., 1992), and approximately one third of the postsynaptic targets of retinotectal terminals are GABAergic (Whyland et al., 2020). GABAergic SC neurons and their connections with other brain regions were shown to play a critical role in wakefulness and eye movements (Mize, 1992; Sooksawate et al., 2012). However, the development and functional organization of GABAergic SC circuits is still largely unknown. Several GABAergic SC cell types have been described based on axon projections, orientation of dendritic arbors, and/or the colocalization of a variety of immunocytochemical markers (Behan et al., 2002; Gale and Murphy, 2014, 2018; Whyland et al., 2020). For example, selective labeling of inhibitory neurons in the sSC of Gad2–Cre mice revealed axon terminals in several nuclei, including the PBg, the dorsolateral portion of dLGN, and the vLGN (Gale and Murphy, 2014). Pax7 is a transcription factor known to direct embryonic cells along a neurogenic lineage and form of superior collicular boundary. In the adult mouse brain, Pax7 expression is particularly high in the sSC (Figures 6A and 6B), and in our snRNAseq data, we found that *Pax7* was expressed strictly in the inhibitory neurons (Figures 6C and 6D). Of the inhibitory neuron subpopulations, about half of them express *Pax7* (i.e. IN2, IN4, IN6, IN9, IN11 and IN12) (Figure 6D). To determine whether Pax7 expression labels an anatomically heterogeneous or homogeneous class of inhibitory SC neurons, we sought to determine their axonal projection. To this end, we injected AAV-FLEX-GFP into the superficial SC of adult Pax7-Cre mice in which GFP expression is limited to the Pax7-expressing cells. Of note, Pax7-Cre mice crossed to Rosa26-Tomato reporter mice showed Tdtomato expression predominantly in the midbrain, and not in the cortex (Figure 6E), consistent with the *Pax7* expression pattern shown in the Allen Brain Atlas (Figure 6A). In the mice injected with AAV-FLEX-GFP, we found GFP^+^ axon terminals in the vLGN but not in the LP, PBg or dLGN (Figures 6F, 6G, and 6H). This result indicates that Pax7 labels a homogeneous population of inhibitory SC neurons based on their axonal projection.

**Figure 6.**
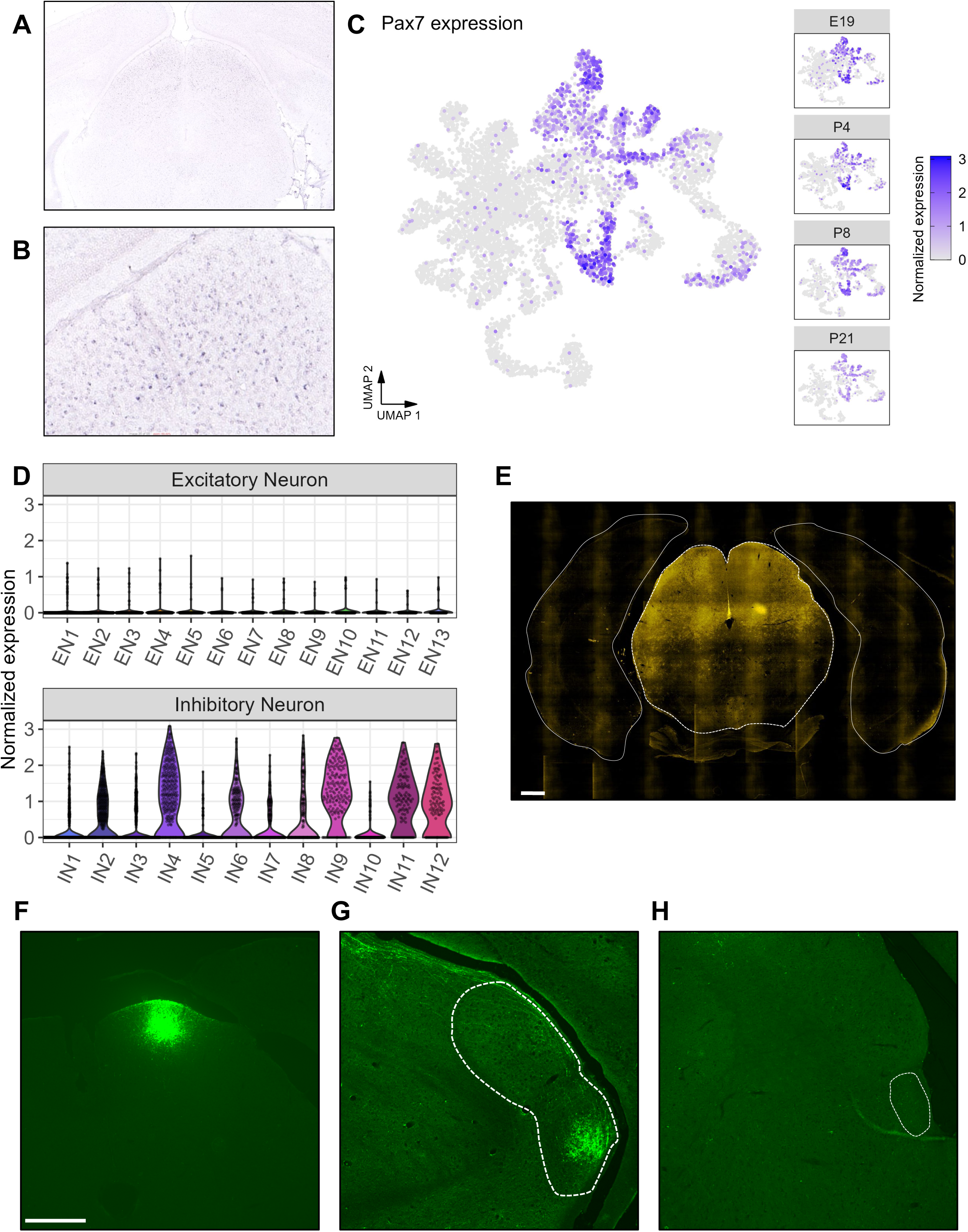
Pax7+ cells are a population of GABAergic neurons with a common downstream brain target. **A**) In situ hybridization against *Pax7* from Allen Brain Atlas. *Pax7* expression is restricted to the midbrain. **B**) High magnification of the boxed area in Figure 6A. **C**) Left: UMAP of *Pax7* expression in SC neurons. Right: UMAP of *Pax7* expression in SC neurons across the four developmental time points collected. **D**) Violin plot of *Pax7* expression in excitatory and inhibitory neuron subtype demonstrating that detection of *Pax7* is limited to a subset of inhibitory neurons. **E**) Tdtomato expression. A coronal section of brain from Pax7-Cre; Rosa26-TdTomato double transgenic mouse. Consistent with the in situ hybridization (Figure 6A), TdTomato expression is restricted to the midbrain. **F-H**) GFP expression in the SC (**F**), LGN (**G**), and PBg (**H**). White dotted areas indicate the respective nuclei. Adult Pax7-Cre; Rosa26-TdTomato received injection of AAV2-FLEX-GFP into the superficial SC. Scale bars, 500 µm.

## DISCUSSION

The superior colliculus is a region of multisensory integration, and a rising model for studying circuit formation. In this study, we describe diverse cell types and neuronal subtypes that make up the mouse SC throughout the distinct stages of brain development. We show that each cell and neuronal subtype is marked by unique sets of genes with many genes exhibiting age-dependent changes in expression. Thus, our study expands previous sc-and snRNAseq studies and identifies cell populations and their unique transcriptional signatures in developing as well as mature SC. To facilitate open exploration of this data, we generated a web portal for analyzing and comparing gene expression profiles in the SC cell types and neuronal subtypes across the ages (https://parklabmiami.shinyapps.io/superior-colliculus-snRNAseq/).

During development, distinct transcription factors, cell adhesion molecules, and extracellular matrix proteins contribute to axon guidance and formation of proper neural circuits. Our study shows dynamic gene expression changes in neurons across different time points. Of those differentially expressed genes, one notable gene is *Zic1.* We found that Zic1 is highly expressed in the deep SC early during development (i.e. E19 and P4) but expression subsides by adulthood. Moreover, expression of *Zic1* is high in the medial but low in the lateral portion of SC, suggesting that Zic1 may play a role in region-specific targeting of axons in SC. It has been long known that members of the Zic family of zinc finger transcription factors regulate the expression of axon guidance molecules and play critical roles in axon targeting in the CNS. For example, Zic2 controls the expression of EphB1 receptors in select populations of RGCs in the retina which in turn promotes projection of these neurons to the ipsilateral brain side (Lee et al., 2008). It will be interesting to examine in the future whether Zic1 plays a role in guiding incoming axons to proper regions in the SC. Another notable gene is *Cdh22,* which we find to be specific to inhibitory neurons. Previous studies have shown that certain cadherin members have a function specifically in interneurons, particularly at the inhibitory synapses; Cdh13 deficiency was shown to reduce synapse turnover, producing an increase in inhibitory synapse density (Rivero et al., 2015). It is plausible that Cdh22 plays a similar role and regulates interneuron development and synaptic function in the SC.

Besides the gene expression changes seen in neurons, we also observed changes in the expression of many guidance molecules in astrocytes. Studies have shown that glial cells play important roles in the formation and maturation of CNS circuits. For example, studies have shown that immature astrocytes help guide axons to their targets by providing appropriate cell surface receptors and adhesion molecules. Astrocytes also secrete a variety of growth factors and extracellular matrix proteins that modulate the direction of axon growth and extent of synapse formation (Eroglu and Barres, 2010). Notably, our data showed dynamic changes in astrocytic expression of various guidance molecules across time points; while genes encoding Eph receptor and Ephrin molecules showed decreased expression over time, subsets of genes from the Semaphorin-Plexin signaling pathway, such as *Sema4a* and *Sema4b,* increased in expression. It remains to be determined whether expression of these genes in astrocytes plays a functional role during SC development.

The SC contains a heterogeneous population of GABAergic neurons whose properties and innervation play a key role in the regulation of collicular functions. We showed that *Pax7* transcripts are enriched in subpopulations of inhibitory neuronal subtypes (i.e. approximately half of all inhibitory neurons in the SC). By labeling axons from Pax7^+^ neurons in the sSC, we found that these neurons send long projections exclusively to vLGN and not to other known distal targets (e.g. LP, dLGN and PBg) (Benavidez et al., 2021; Gale and Murphy, 2014). The vLGN is thought to be a part of a multimodal system, receiving retinal afferents but establishing subcortical interconnections with non-imaging forming areas (e.g. olivary pretectal nuclei, accessory optic system, and suprachiasmatic nuclei) (Swanson et al., 1974). The vLGN also projects to the SC, the periaqueductal gray area (PAG), and the nucleus reuniens of the ventral midline thalamus. These regions all contribute to behavioral responses to visual threats (Moore et al., 2000; Salay and Huberman, 2021). Whether Pax7^+^ sSC neurons are a functionally distinctive population of cells that subserve visually guided behavior or non-image-forming visual functions remains unknown.

In summary, this report provides a comprehensive molecular characterization of the cell types that comprise the SC across murine development, with particular emphasis on neuronal heterogeneity and molecules involved in circuit assembly. By comparing *in situ* hybridization data from the Allen Brain atlas, we demonstrate that certain transcriptionally defined neuronal subtypes also display anatomic specificity. We also attempt to provide a unified definition of neuronal subtypes of the SC across multiple studies and to infer subtype-specific projections to various brain regions. Finally, by leveraging our snRNAseq data, we confirm and characterize projections from the Pax7^+^ subset of SC neurons to demonstrate the utility of this data as a starting point for manipulations in specific neuronal populations and circuits.

## Acknowledgments

This work was supported by grants from the National Eye Institute (NEI) R01EY022961 (K.K.P.), NEI R01EY032542 (K.K.P), NEI 1U01EY027257 (K.K.P), NEI R21EY031026 (K.K.P), DOD W81XWH-19-1-0736 (K.K.P), The Miami Project to Cure Paralysis, and the Buoniconti Fund (K.K.P) and Glaucoma Research Foundation (K.K.P). We thank Dr. April Mann for manuscript editing.

## Author Contributions

A.C.A., J.S.C., and K.K.P. designed the experiments, conducted the experiments and wrote the paper. R.M., and K.L. assisted the experiment. F. B. contributed to data analysis.

## METHODS

### Mice

All animal experimental procedures were performed in compliance with protocols approved by the Institutional Animal Care and Use Committee (IACUC) at the University of Miami. Animals used were C57BL/6J (The Jackson Laboratory, stock# 000664), Pax7-Cre (The Jackson Laboratory, stock# 010530), and R26 loxP-STOP-loxP-tdTomato (Arenkiel et al., 2011)(a gift from Dr. Fan Wang, Massachusetts Institute of Technology). All animals were housed in a viral antigen-free facility and kept under standard 12-h light-dark conditions. For all surgical procedures, mice were anesthetized with ketamine and xylazine. For analgesia, buprenorphine (0.05 mg/kg) was administered post-operatively. Animals of both sexes were used unless specified.

### Nuclei isolation and analysis of single-nucleus RNAseq

Nuclei isolation was performed following the protocol described previously (Velmeshev et al., 2019). Briefly, SC from E19, P4, P8, and P21 mice were isolated and homogenized in RNAase-free lysis buffer (0.32 M sucrose, 3 mM CaCl^2^, 3 mM MgAc^2^, 0.1 mM EDTA, 10 mM Tris-HCl,1 mM DTT, 0.1% Triton X-100 in DEPC-treated water) using a glass dounce homogenizer on ice. Animals of both sexes were used for all ages except for E19 which were all males. The homogenate was loaded into a polycarbonate ultracentrifuge tube containing sucrose solution (1.8 M sucrose, 3 mM MgAc2, 1 mM DTT, 10 mM Tris-HCl in DEPC-treated water) in the bottom and centrifuged at 107,000 g for 2.5 hours at 4°C. Supernatant was aspirated, and the nuclei containing pellet was incubated in RNAse-free 1x PBS, 0.04% BSA, 0.2 U/µl RNAse inhibitor on ice before resuspending the pellet. The nuclear suspension was filtered twice through a 30 µm cell strainer and counted using a Nexcelom Cellometer K2 before performing single-nucleus capture on the 10X Genomics 3’ v3 single cell RNA-Seq. Target capture of 2,000 nuclei per sample was used and the 10X 3’ v3 scRNA-Seq library preparation performed and sequenced on the NovaSeq SP 100 (200,000 reads/nucleus) by the Oncogenomics Core Facility at University of Miami Miller School of Medicine.

After sequencing, Illumina output was processed using CellRanger v3.0.2. Base call files for each sample were demultiplexed. A pre-mRNA reference was generated with CellRanger mkref using the mm10 mouse genome. Each sample was aligned to the custom mm10 mouse reference genome using CellRanger. Sample reads were sequenced across two lanes and concatenated after alignment, resulting in a single count matrix per sample.

#### Pre-processing and quality control

To distinguish nuclei-containing droplets from empty droplets, we performed cell calling on the unfiltered UMI count matrices using a combination of barcode-ranking and the empty-droplet detection `emptyDrops` function as implemented in the `DropletUtils` R package (Lun et al., 2019). First, nuclei were ranked according to total UMI count and visualized in a log-total UMI vs log-rank plot (Figure S1A). A spline curve was fit to the data to identify “knee” and inflection points, and cells with total UMI count above the knee were considered nuclei-containing droplets. Next, we used the `emptyDrops` algorithm to further distinguish empty droplets from nuclei for nuclei with lower total UMI counts.

To remove potential doublets, we applied the Python package `Scrublet` (Wolock et al., 2019) to each individual sample using default parameters. In brief, `Scrublet` simulates multiplets by sampling from the data and builds a nearest-neighbor-based classifier. Cells with high doublet scores were flagged and removed.

We performed further quality control based on metrics such as total UMI, number of unique genes detected, and mitochondrial transcript content (Figure S1B). Lower-bound thresholds for total UMI and unique gene detection rates were determined by computing three absolute median deviations (MADs) below the median. Across all samples, mean total UMI was 10,556 (E19, 7,457; P4, 9,882; P8, 10,957; P21, 13,928), mean total genes detected was 3,550 (E19, 3,030; P4, 3,574; P8, 3643; P21, 3,953), and mean mitochondrial transcript content was 0.62% (E19, 0.70%; P4, 0.98%; P8, 0.41%; P21, 0.39%). Remaining high-quality nuclei were used for downstream analysis.

#### Integrated identification of all cell types across development

We first performed standard single-nucleus RNAseq analysis using the Seurat R package (v4.2.1)(Stuart et al., 2019). We observed significant batch effects between samples from each developmental time point; for example, neurons from P21 clustered separately from neurons from all other time points due to detection of contaminant ambient RNAs such as *Plp1* (Figure S1C). Based on this observation, we determined that Data Integration was necessary.

To better identify shared and unique cell types across all developmental time points, we performed integrated analysis as outlined in Seurat’s Data Integration workflow (Butler et al., 2018). In brief, after UMI count matrices were log-normalized, the top 2,500 variable genes and first 15 principal components were used for dimensional reduction and clustering. Cell types were identified using a combination of differential expression testing, comparisons against reference data sets, and prior knowledge of cell type-specific marker genes. For differential expression testing to identify markers, we used the `FindAllMarkers` function in Seurat using default parameters. For comparisons against reference data sets, we used the `SingleR` R package (Aran et al., 2019). See methods section on “Comparison of neuronal subtypes to reference data” for description of the `SingleR` method. After cell type identification, differentially expressed genes were recomputed again using the `FindAllMarkers` function. We principally used the following genes for cell type identification: Excitatory neurons, *Slc17a6*; Inhibitory neurons, *Gad1*; Astrocytes, *Aqp4* and *Gfap*; Oligodendrocyte-lineage cells, *Cspg4*, *Bmp4*, *Mbp*; Dividing cells, *Mki67*; Microglia, *P2ry12*; Endothelial cells, *Cldn5*; Vascular Leptomeningeal cells, *Col1a1*; Epithelial cells, *Cdh1*.

#### Integrated analysis of neurons across development

To investigate neuronal heterogeneity and identify neuronal sub-types, we performed a similar integrated analysis as described above with certain modifications. First, we used the top 300 variable genes and top 10 principal components for dimensional reduction and clustering. Principal components were determined using the “elbow” plot heuristic. We also set the “resolution” parameter in the `FindClusters` function to 0.72 based on empirical observations that at this resolution neuron clusters could be identified using single or a small set of genes. Neuronal subtypes were annotated using these parameters. Neuronal subtype marker genes were computed using the `FindAllMarkers` function and filtered by taking the top 2 genes by p-value and then by log(fold-change). To better identify subtype markers within each of the excitatory and inhibitory neuron classes, we extracted each class and reperformed differential expression tests via `FindAllMarkers`. We used the default Seurat log2(fold-change) threshold of 0.25 and adjusted p-value threshold of 0.05 for determining the number of DEGs (Figures S2E and S2F). We further quantified the similarity between neuron subtypes by constructing a dendrogram relating the average expression profile of neuron subtypes using the same genes used for cluster analysis (Figure S2D).

We also sought to identify gene expression changes in all excitatory or inhibitory neurons across development using the `FindMarkers` function. We observed that many of the top differentially expressed genes between time points were non-neuronal, e.g the gene *Ttr*, which has been shown to be specific to cells of the choroid plexus in the CNS (Herbert et al., 1986). We reasoned that many of these genes may be derived from the ambient RNA during droplet processing (Caglayan et al., 2022). To mitigate the effects of ambient RNA contamination in differential expression testing, we applied ambient profile estimation algorithms in the `DropletUtils` R package (Figure S1A). In brief, unfiltered UMI count matrices, containing transcript quantifications for all 10X Chromium lipid encapsulations, were used to estimate the ambient RNA profile based on expression data from low total UMI droplets. We used the `ambientContribMaximum` function to then filter out genes from differential expression test results which had an average maximum ambient RNA count contribution greater than 20% across all samples in the comparison. This approach was also applied in identifying gene expression changes across developmental time points for non-neuronal populations.

To further characterize the global changes in neuronal gene expression across development, we combined all neuronal cells into a single group and performed time point comparisons as well as ambient RNA filtering (Supplementary Figure S3A). We performed Gene Ontology enrichment analysis for Biological Process terms using these DEGs using the `topGO` R package (Figure S3B) (Alexa A, Rahnenfuhrer J (2022). topGO: Enrichment Analysis for Gene Ontology. R package version 2.50.0).

#### Comparison of neuronal subtypes to reference data

To compare the neuronal subtypes identified in our study to neuronal subtypes described previously in other reports studying SC neurons, we applied the `SingleR` algorithm from the `SingleR` R package. In brief, `SingleR` first identifies marker genes for each neuron subtype label in the reference data in a pair-wise manner. These genes are then used to compute Spearman correlations between the gene expression profiles of cells from the query dataset and cells of the neuron subtypes from the reference dataset. For each query cell, a per-subtype distribution is generated from the correlation values against cells from that neuron subtype label. For that query cell, the per-subtype label is defined as a fixed quantile (default 0.8) of this distribution. The subtype label with the highest score becomes the predicted subtype of the query cell. Heatmaps demonstrating the per-reference-label contribution to each neuronal subtype in the current study were generated using these labels.

#### Investigation of adhesion and axon guidance molecules

To perform a comprehensive query of cell adhesion and axon guidance molecules, we pulled gene sets from the following sources: extracellular matrix and adhesion molecules were pulled from GeneCopoeia’s ExProfile^TM^ Extracellular Matrix and Adhesion Molecules gene panel; axon guidance molecules were pulled from the KEGG pathway database using the pathway ID “mmu04360”. To investigate the heterogeneity of expression of these molecules in SC nuclei, we performed integrated cluster analysis using the Seurat integration pipeline as described above using these gene sets. We further performed Principal Component Analysis (Figure S5A) and Multidimensional Scaling analysis (Figure S5B) to quantify cell type heterogeneity based on these genes.

### Assessment of projections of Pax7-expressing SC neurons

For Cre-dependent anterograde labeling of Pax7 expressing neurons in the superficial SC (sSC), approximately 100 nl of AAV2–CAG–FLEX–GFP (University of North Carolina Vector Core) was injected into the sSC of Pax7-Cre mice (10 weeks old). Injection coordinates were as follows: posterior from bregma, lateral from midline, and depth in mm, 4.16, 0.2, 0.5, and 1.0-1.2, respectively. Three to 4 weeks after AAV injection, mice were anesthetized and then transcardially perfused with 4% paraformaldehyde in phosphate buffered saline (PBS). Brains were dissected and postfixed in 4% paraformaldehyde in PBS for 16 hours, and cryoprotected in 30% sucrose in PBS for 2-3 days. Brains were embedded in OCT compound (Tissue-Tek) and coronal sections (20 μm) were cut using a cryostat. Sections were immunostained by incubating in primary antibodies in 5% Normal Goat Serum in PBS with 0.3% Triton-X overnight at 4 °C. Primary antibodies used were: RFP (Rockland 600–401-379S, 1:1000) and GFP (Abcam ab13970, 1:2000). Following primary antibody incubation, sections were washed and incubated in species-appropriate Alexa Fluor IgG (H + L) secondary antibodies (Invitrogen, 1:500) at room temperature for 1 hour. Slides were mounted using Vectashield with DAPI (Vector Laboratories H-1200). Images were obtained using a Nikon Eclipse Ti fluorescent microscope or an Olympus FluoView 1000 confocal microscope.

#### Fluorescent in situ hybridization (FISH)

RNAscope FISH was performed on 20 μm thickness coronal brain sections from adult mice (8 weeks old) using the RNAscope® Multiplex Fluorescent v2 Assay (ACD Biotechne, Catalog No. 323100) according to the manufacture’s protocol. Target probes used are as follows: RNAscope® Probe - Mm-Gfra1 (Cat No. 431781), RNAscope® Probe -Mm-Slc17a6-C2 (Cat No. 319171-C2). TSA-based fluorophores were from Perkin Elmer (TSA Plus Fluorescein, PN NEL741001KT; TSA Plus Cyanine 3, PN NEL744001KT; TSA Plus Cyanine 5, NEL745001KT). Images were acquired using an Olympus Confocal FV1000 microscope or an Andor Dragonfly confocal microscope.

#### Data Availability

All newly generated raw sequencing reads and key processed files are available at Gene Expression Omnibus (GEO) under the accession GSE224407. Processed data from the Trans-seq study (Tsai *et al*. 2022) were pulled from links from the Github repository associated with the publication (https://github.com/duanxlab/Trans-Seq). Trans-seq count matrix data were accessed from GEO with the accession GSE202257. Processed data from the Vector-seq study (Cheung *et al*. 2021) were pulled from links from the Github repository associated with the publication (https://github.com/vic-cheung/vectorseq). Vector-seq count matrix data were accessed from GEO with the accession GSE189907.

#### Code Availability

The code to analyze snRNA-seq data is available in Github (https://github.com/ParkLabMiami/snRNAseq-developing-superior-colliculus).

## Supplementary Figures

**Figure S1.**
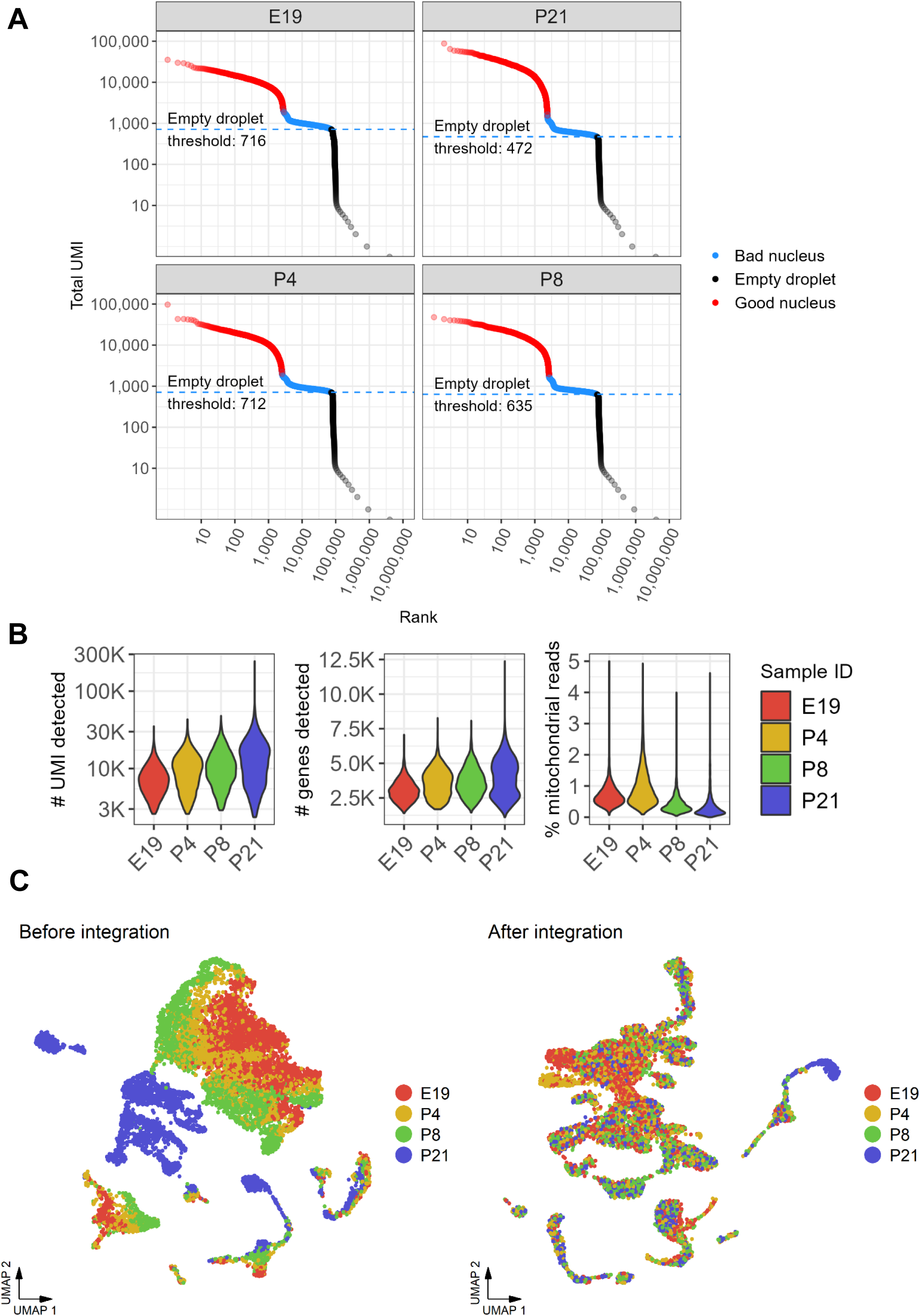
snRNAseq preprocessing, quality control, and batch correction. **A**) Barcode-rank plots for each sample. Droplets containing 10X Genomics GEM barcodes are ranked by their total UMI count. Dots colored in red indicate high quality nuclei. Dots colored in blue indicate putative low-quality nuclei. Dots in black are presumed empty droplets containing no nuclei (see Methods for more details on use of `emptyDrops()` function from `DropletUtils` R package). **B**) Violin plots of various quality control metrics used for additional filtering of low-quality nuclei. **C**) UMAP plots of nuclei before (left) and after (right) batch correction implemented through Seurat’s Data Integration method. Note that neuronal nuclei from P21 (left UMAP, blue clusters in center-upper-left) do not cluster with other neuronal nuclei from other samples (red, yellow, and green cluster in upper-right).

**Figure S2.**
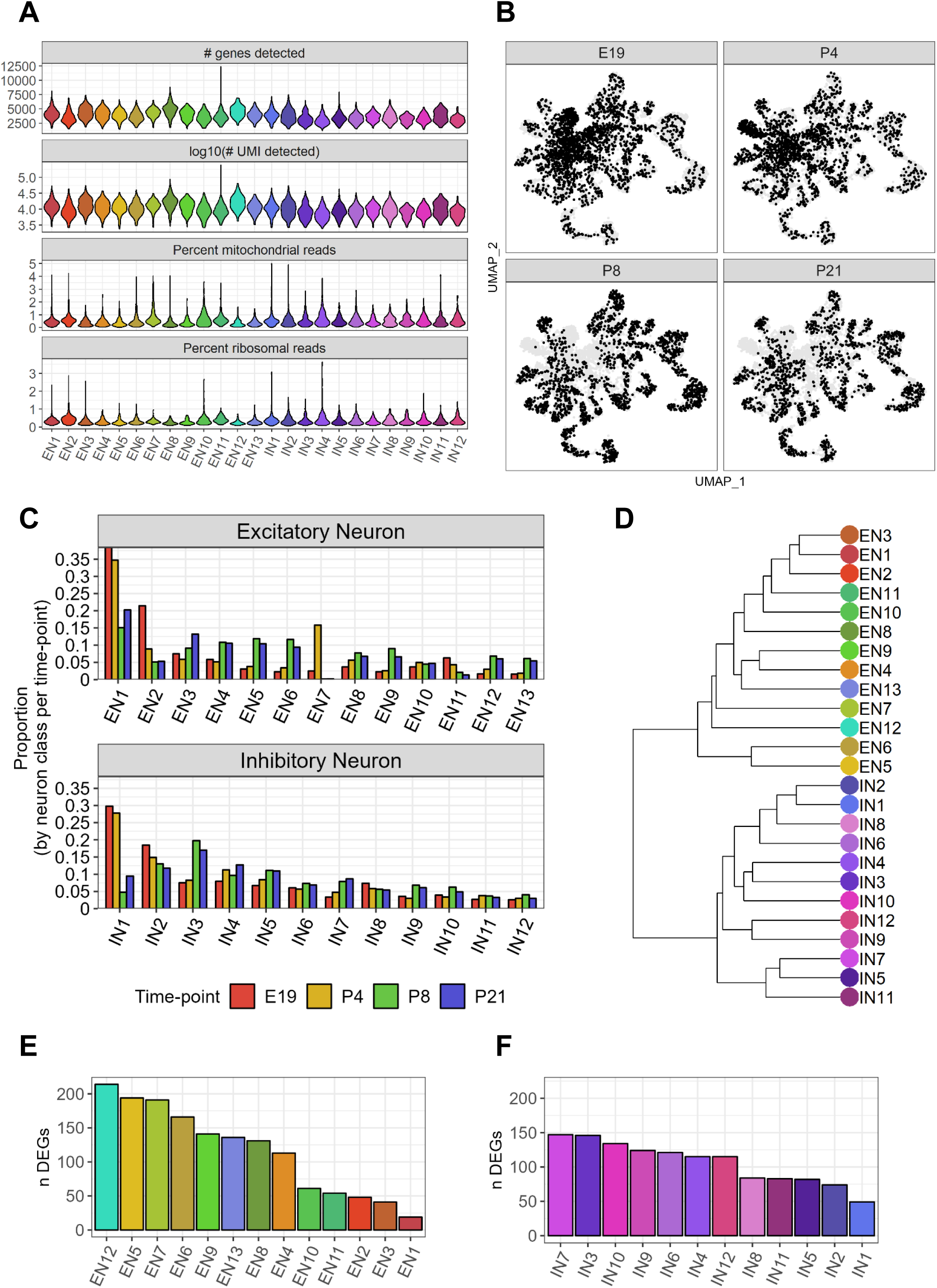
Cluster analysis of excitatory and inhibitory neurons. **A**) Various quality control metrics per neuronal subtypes indicating that nuclei data quality was not a driving factor in subtype analysis. **B**) UMAP of neuronal nuclei shaded by developmental time point. Note the gradual decrease of neurons located at the center of the UMAP (primarily EN1 and IN1) and increase in distally positioned neurons. **C**) Quantification of neuronal subtypes as a proportion of all neurons per developmental time point e.g. EN1 is approximately 35% of all excitatory neurons at E19, and IN5 is approximately 10% of all inhibitory neurons in P8. **D**) Dendrogram of neuronal subtypes quantifying the relationship between neuronal subtypes. **E**) Quantification of the number of differentially expressed genes distinguishing each excitatory neuronal subtype against all other excitatory neurons. **F**) Similar quantification as (E) but for inhibitory neuron subtypes. Differential expression computed using Wilcoxon Rank Sum test as implemented by the `FindMarkers()` function in Seurat R package.

**Figure S3.**
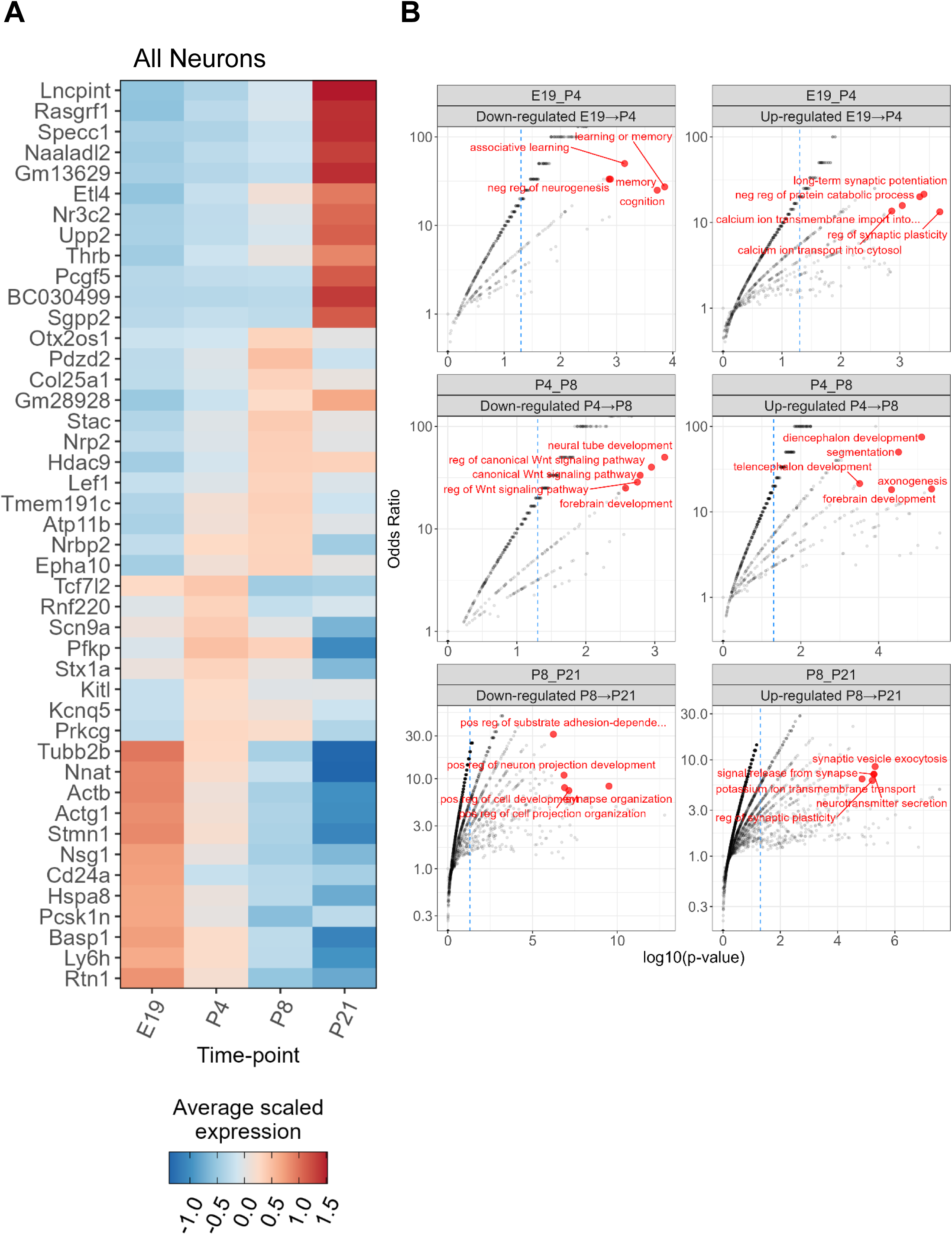
Pan-neuronal gene expression changes. **A**) Heatmap of differentially expressed genes between developmental time points where all neuronal subtypes are first grouped before computing genes enriched per time point compared against all other time points combined. **B**) Gene Ontology:Biological Process enrichment analysis using differentially expressed genes as computed in (A) but split between up-and down-regulated genes between sequential time points. Y-axis denotes the odds ratio between the number of significant differentially expressed genes and the total number of genes annotated to each biological process. The top 5 terms per comparison with at least 3 significant genes are highlighted in red. Dashed blue line corresponds to a p-value cutoff of 0.05. Differential expression computed using Wilcoxon Rank Sum test as implemented by the `FindMarkers()` function in Seurat R package. GO analysis performed using the “classic” algorithm for the Fisher test as implemented in the `topGO` R package.

**Figure S4.**
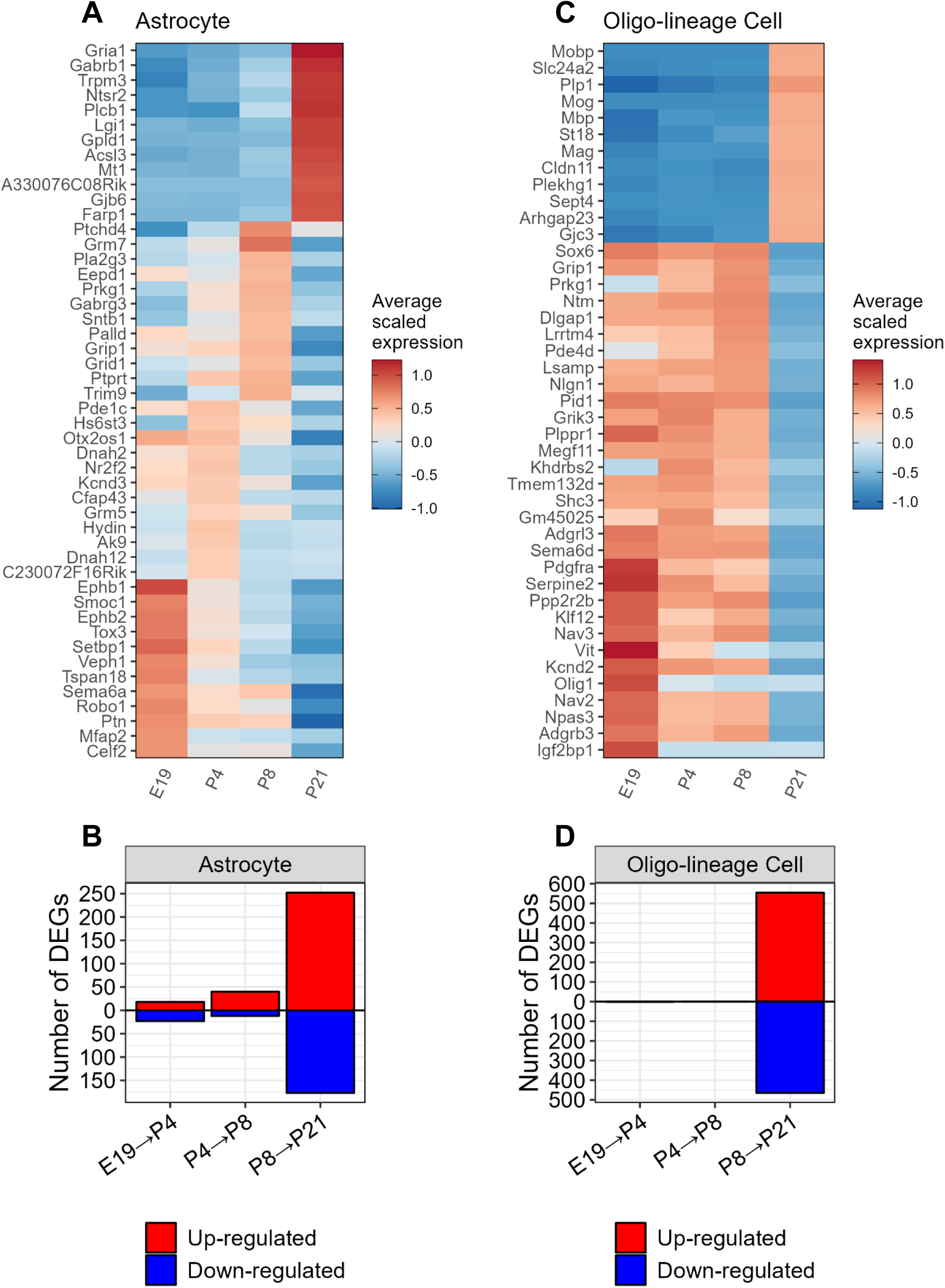
Gene expression changes in astrocytes and oligodendrocyte-lineage cells. **A**) Heatmap of differentially expressed genes in astrocytes where astrocytes at each time point are compared against all other time points combined. **B**) Quantification of the number of differentially expressed genes in astrocytes when comparing sequential developmental time points. **C**) Heatmap of differentially expressed genes in oligodendrocyte-lineage cells, similar to (A). **D**) Quantification of number of differentially expressed genes in oligodendrocyte-lineage cells, similar to (B).

**Figure S5.**
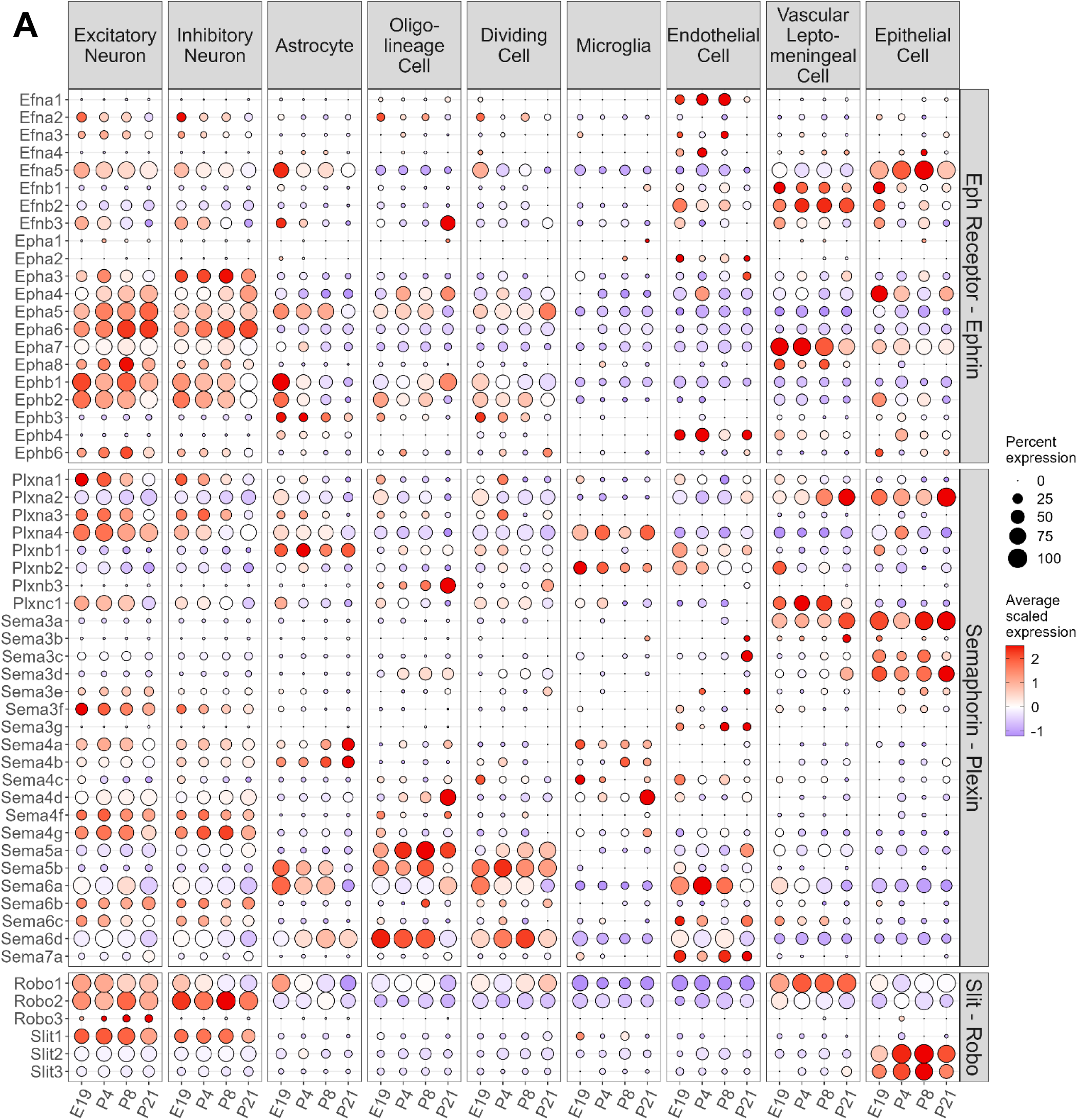
Comprehensive panel of axon guidance molecules reveals cell type-specific expression. Dot plot of axon guidance molecule gene expression by cell type. Genes are grouped by molecule family on right y-axis. Expression values represent the averaged scaled expression values of nuclei grouped by cell type and time point.

**Figure S6.**
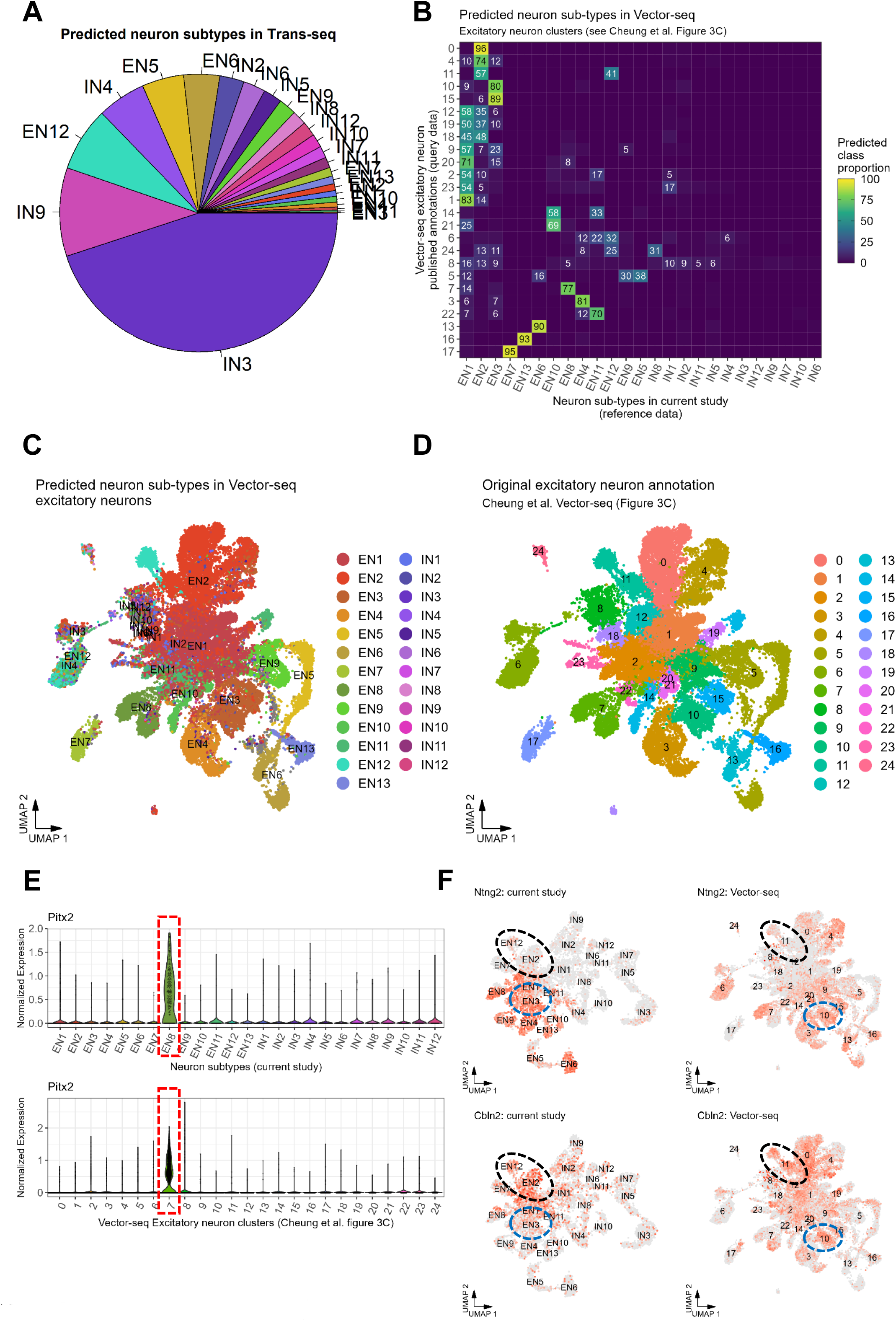
Extended `SingleR` analyses comparing our dataset to Trans-seq and Vector-seq datasets. **A**) Pie graph of the composition of predicted neuronal subtypes that comprise the entire Trans-seq dataset. **B**) Heatmap of predicted subtype proportions that comprise each of the excitatory neuron clusters as defined in Vector-seq (see Cheung *et al*. figure 3C). **C** and **D**) UMAP of predicted subtype labels (**C**) compared to the original cluster labels (**D**) of *Slc7a6*^+^ excitatory neurons from Vector-seq. UMAP coordinates are identical to figure 3C from Cheung *et al*. **E**) Violin plots depicting normalized expression *Pitx2* in neuron subtypes of the current study (top) or excitatory neuron clusters from Vector-seq (bottom). Dashed red circle indicates predicted corresponding clusters between the two studies based on `SingleR` results. **F**) UMAP depicting expression of *Ntng2* (top row) and *Cbln2* (bottom) for neuron subtypes in current study (left column) or excitatory neuron clusters from Vector-seq (right column). Dashed black and blue ovals indicate corresponding clusters between the two studies based on `SingleR` results.

